# Cognitive Endophenotypes of Modern and Extinct Hominins Associated with *NTNG* Gene Paralogs

**DOI:** 10.1101/034413

**Authors:** Pavel Prosselkov, Ryota Hashimoto, Denis Polygalov, Kazutaka Ohi, Qi Zhang, Thomas J. McHugh, Masatoshi Takeda, Shigeyoshi Itohara

## Abstract

A pair of vertebrate-specific and brain-expressed pre-synaptic genes, *NTNG1* and *NTNG2,* contributes to the Intellectual Quotient (IQ) test scores in a complementary manner. Single nucleotide polymorphisms (SNPs) of *NTNG1* are associated with attenuated verbal comprehension (VC) or processing speed (PS) while *NTNG2* SNPs affect working memory (WM) and perceptual organization (PO), forming cognitive endophenotypes in healthy and schizophrenia (SCZ)-affected human subjects. Regions of interest (ROIs), defined as 21 nucleotide (nu) long *NTNG* gene loci symmetrically embedding the IQ-affecting mutation alleles (VC/PS and WM/PO), underwent dramatic evolutionary changes from mice through primates to hominins, at the accelerated rates. Mutation alleles associated with the higher VC and WM IQ scores are found in the genomes of extinct hominins of Neolithic times, however, lower WM scores associated allele is also detectable in Mesolithic hunters genomes. Protein sequence of *NTNG1* is 100% conserved among the primates, archaic and modern extinct hominins while *NTNG2* underwent a recent selection sweep encoding a primate-specific S371A/V (~50,000 yrs BC), and a modern human (5,300 yrs BC) T346A substitutions. We show that a 500 mln yrs old genomic duplication of a synapse primordial gene provided a substrate for further synapse elaborations and its ultimate capacitive expansion of what evolved into a vertebrate cognitive superior complexity – intelligence.

## INTRODUCTION

S.Ohno ingeniously proposed that new gene function can result from a gene duplication and following it gene paralogs SF (1). Two gene paralogs, *NTNG1* and *NTNG2,* expressed predominantly in the brain (2), and encoding Netrin-G1 and Netrin-G2 proteins, respectively, are expressed pre-synaptically and segregate in a non-overlapping manner into distinct neuronal circuits (3, 4). They are related to the netrin family of axonal guidance cues (5) but differ in that they attach to the axonal membrane via a glycosylphosphatidylinositol (GPI) link (6, 7), a known lipid raft-associated membrane signaling cascade organiser (6, 8). The first evolutionary precursor of *NTNG* as a single gene copy can be located in the genome of a primitive vertebrate tunicate/urochordate *Ciona intestinalis* (sea squirt, ENSCING00000024925), reported to be the first organism with the neural crest primordials (9), multipotent brain progenitor cells (10), and neurogenic placodes, facilitating the transition from pelagic invertebrate life style to a predatory vertebrate (11). The dramatic expansion of human cerebral cortex over the course of evolution (12–15) had provided new niches for accommodating either *de novo* or advancing pre-existed cognitive features and culminating in the positively selected human cognitive functions (16).

IQ tests are a surrogate measure of general human cognitive ability characterising intelligence. They are often administered as WAIS-III/IV (17) and represent a cumulative score of 4 cognitive indices: VC, WM, PO and PS frequently referred as “cognitive domains” (18). IQ has been validated by factor analyses (19), and a common factor (correlate) influencing each of them frequently referred as g, or “general intelligence”, proposed by Spearman (20), but recently challenged as the only existing correlate (21). IQ is affected by several mental disorders including schizophrenia (SCZ) characterised by severe cognitive deficits in WM (22), behavioral flexibility and resulting in low performance in cognitive tests (23–25). Since both *NTNG* paralogs have been reportedly associated with SCZ (26–34) we investigated whether these gene paralogs contribute to human intelligence by assessing the IQ of human carriers for the *NTNG1* and *NTNG2* SNP alleles against non-carriers with and without SCZ.

## RESULTS

Several *NTNG1* and *NTNG2* SNPs have been previously reported to be associated with SCZ (27, 30). We have found that out of 11 SNPs tested, five affect the IQ scores and composite domains in human subjects (Figure 1, Supplementary Table 1 (ST1)). SCZ patients carrying rs2218404 T allele (Figure 1A–1) of *NTNG1* (T/G and T/T genotypes, N = 25 patients) compared with G/G genotype (N = 36 patients) demonstrated attenuated full-scale IQ (FIQ, ANCOVA*p* = 0.0057 (F = 7.80)), and VIQ (*p* = 0.0033 (F = 8.87)). VC domain score was the main contributor to the VIQ decline (*p* = 0.0050 (F = 8.08), Figure 1B–1), with low scores across all parameters except comprehension (CH): vocabulary (Vc, *p* = 0.020 (F = 5.49)), similarities (SiM, *p* = 0.041 (F = 4.23)), and information (IF, *p* = 0.0067 (F = 7.50), Figure 1B–1 lower panel). Thus, a mutation allele of *NTNG1,* rs2218404, is associated with low VC and affects the VIQ via the attenuated Vc, SiM and IF subscores. The next *NTNG1* SNP found to affect IQ was rs96501, with attenuated PS in healthy human subjects C allele carriers (N = 45) vs T/T genotypes (N = 98, *p* = 0.028 (F = 4.89), Figure 1B–1) with no effect on SCZ patients. The contributing affecting score was symbol search (SS, *p* = 0.053 (F = 3.79), Figure 1B–1, lower panel) with digit symbol coding (DSC) being also attenuated but non-significantly (*p* = 0.12 (F = 2.40)). Three other SNPs mapped to *NTNG2* (Figure 1A–2) have been also shown to affect IQ. Healthy carriers of the *NTNG2* SNP rs1105684 A allele (N = 49) showed a lower FIQ (*p* = 0.018 (F = 5.70)), VIQ (*p* = 0.029 (F = 4.90)), and PIQ (*p* = 0.048 (F = 3.99)) when compared with the T/T genotypes (N = 96, Figure 1B–2). To check for a potential dosage-dependent effect of a mutation allele on IQ score, *NTNG2* SNP rs2149171 SCZ and healthy human subject cohorts were each split on 3 genotypes, respectively: C/C (N = 14 and 39), C/T (N = 29 and 73), and T/T (N = 15 and 30). The presence of the C allele as a single copy (C/T genotype) was strongly associated with a prominent attenuation in the IQ scores of SCZ patients and was essentially identical to that produced by the C/C genotype when both are compared to the T allele carriers (FIQ: *p* = 0.014 (F = 4.35); VIQ: *p* = 0.029 (F = 3.60); PIQ: *p* = 0.035 (F = 3.42), Figure 1B–2). If in the case of *NTNG1* located SNP rs2218404 the lower VIQ score was contributed mainly by the decreased VC domain scores for Vc, SiM, and IF (Figure 1B–1), in the case of *NTNG2* located rs2149171 the CH and WM domain scores were responsible for the VIQ decline in C allele carriers (*p* = 0.012 (F = 4.54) for CH and *p* = 0.040 (F = 3.27) for WM (N = 12 (C/C); N = 25 (C/T) and N = 14 (T/T)). Similarly to VIQ, where CH of rs2149171 complements the cognitive endophenotype produced by the T-allele of rs2218404, the PIQ attenuated score in the case of rs2149171 was due to lower PO score (*p* = 0.050 (F = 3.04), Figure 1B–2) in the C allele carriers. The third *NTNG2* located SNP found to affect the human IQ was rs2274855 (Figure 1A–2) with a cognitive endophenotype associated with the A-allele presence in SCZ patients (N = 33) vs G/G genotypes (N = 26) and resembling that of the described above C-allele of rs2149171. Accordingly, the attenuated scores were: FIQ (*p* = 0.012 (F = 6.44)), VIQ (*p* = 0.018 (F = 5.70)), and PIQ (*p* = 0.036 (F = 4.46), Figure 1B–2). Similarly to rs2149171 the lower VIQ score was due to declined CH (*p* = 0.035 (F = 4.49)) but unaffected VC that is contrary (complementary) to the rs2218404 endophenotype (Figure 1B–1). WM was robustly affected by the A-allele presence (N = 29; *p* = 0.023 (F = 5.28)) contributed by the low DS score (*p* = 0.026 (F = 5.04)) with LNS and AM being unaffected (Figure 1B–2, lower panel). The observed PO score was comprised by the declined matrix reasoning (MR, *p* = 0.038 (F = 4.34)), block design (BD, *p* = 0.041 (F = 4.23)) with picture completion being unchanged (Figure 1B–2, lower panel). Thus, all three aforementioned *NTNG2* SNPs affect the VIQ and PIQ in human subjects with the first one contributed by the CH subscore and WM and the latter by the PO score (Figure 1C). Contrary to this, the *NTNG1* located SNP (rs2218404) affects the VIQ through the lower Vc, SiM and IF scores and affecting the VC domain scores. Another SNP, rs96501, affects PS domain, although in healthy subjects only.

**Figure 1.**
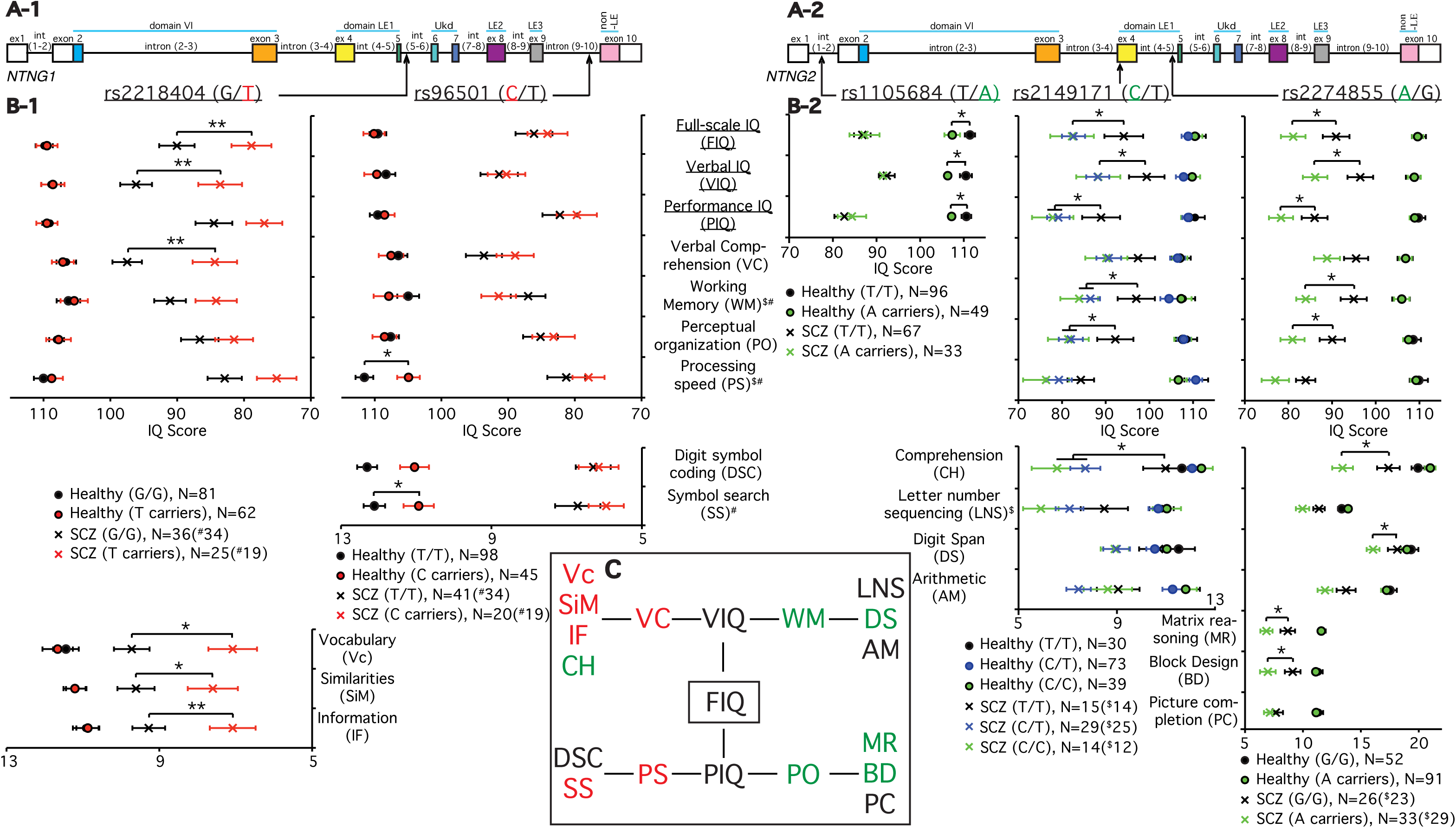
**Complementary effect of *NTNG* paralog SNPs on IQ cognitive domains of human subjects as measured by WAIS-III. (A-1, A-2)** *NTNG1* and *NTNG2* gene structures with the SNPs location indicated. **(B-1, B-2, C)** Affected cognitive domains and scores. Red highlights *NTNG1* and green (or blue, in case of heterozygosity) – *NTNG2* located alleles associated with the attenuated IQ scores. The data are presented as a mean+SEM. *p<0.05,**p<0.01 (two ways ANCOVA (sex, education, and age as co-variates)). The number of human subjects is indicated as N. For statistical and diagnosis details refer to **ST1**.

**Figure 2.**
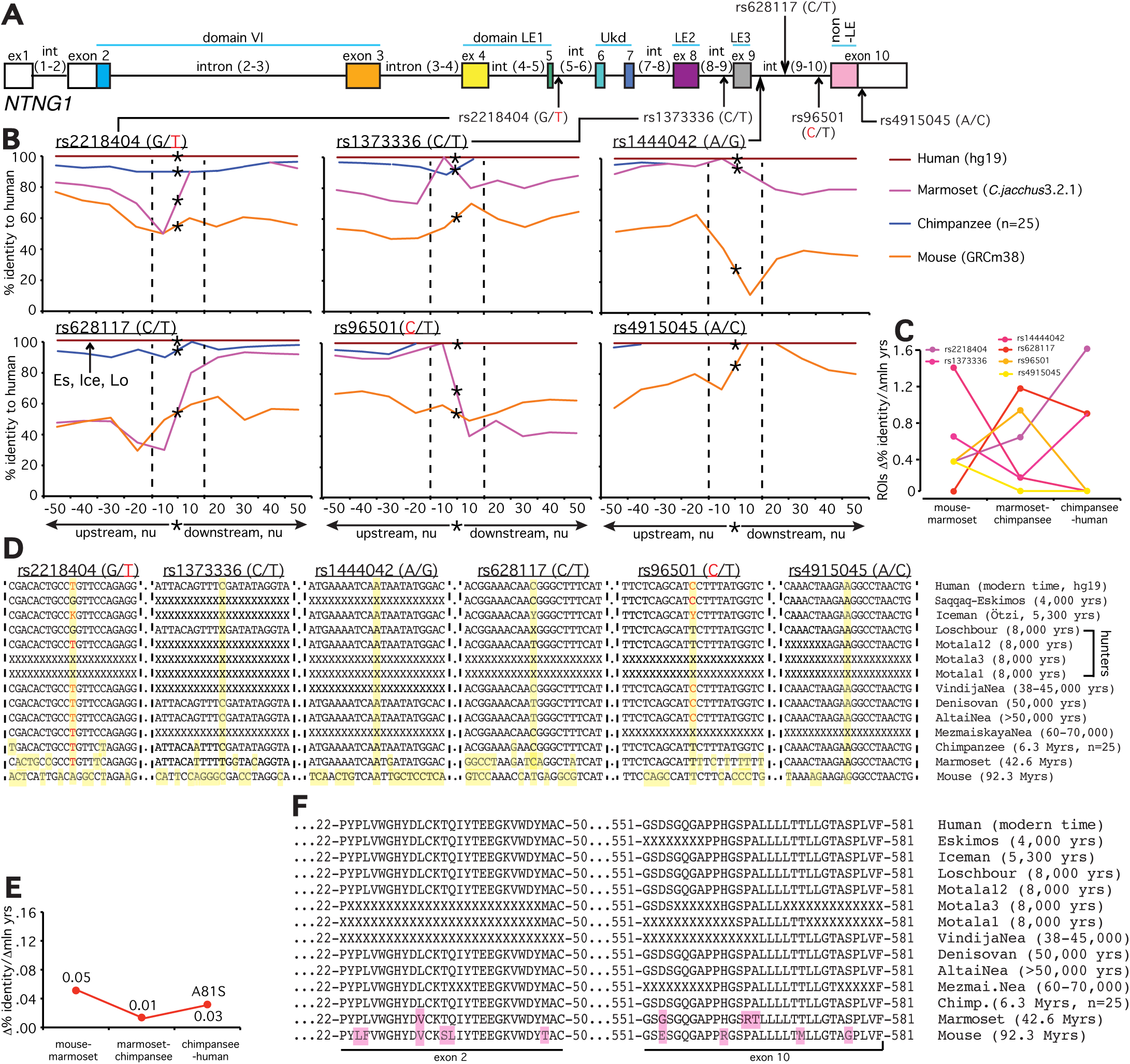
**Accelerated evolution (AE) and definition of a ROI of *NTNG* loci embedding associated with the cognitive endophenotype alleles. Hominins and primate-specific protein amino acid substitutions**. (**A**) *NTNG7* and *NTNG2* SNPs’ gene locations. Alleles associated with the lower IQ scores are shown either in red (*NTNG 7)* or green (*NTNG2).* (**B**) Calculated identity percent of primates and mice to human gene loci as a function of distance from the position of a mutated allele (denoted as a star). Comparison was done in a stepwise manner as +10 nu to the maximum of +50 nu using Stretcher (http://www.ebi.ac.uk/Tools/psa(**70**)). The areas were compared based on the positioning relative to the point mutation without any manual curation for “the best-fit” alignment, but as per the algorithm output only. Due to low level of identity for mouse and marmoset the initial search of the mutation allele positioning was done aligning against the human corresponding full-length intron, and then second time against the obtained 101 nu query (“-50 nu-SNP +50 nu”). Two dashed lines define an area of −10 nu to +10 nu from the mutation allele position. This area (21 nu in total) is defined as a ROI of the given mutation allele of a representing SNP. An arrow indicates a position of extinct hominins-specific mutations (see below). (**C**) Evolutionary rates for the ROIs calculated as a percent identity change relative to the hg 1 9 over the mln of years of evolution. The spectrum color reflects the mutations’ positioning order on a gene, as purple-yellow for *NTNG7,* and yellow-blue for *NTNG2.* (**D**) ROIs DNA sequences across hominins, primates and mice loci. The extinct and ancient hominin’s *NTNG* paralogs were reconstructed from the available datasets (see **SM**): Saqqaq-Eskimos (Es: **59**); Iceman (Ice: **58**); Hunters (Loschbour (Lo), Motala12 (Mo12), Motala3 (Mo3), Motala1: **57**); VindijaNea (Vi: **72**); Denisovan (De), AltaiNea (Al) and MezmaiskayaNea (Me): **73**; chimpanzee (Chi: reconstructed from (71), n=25 animals). Non-available sequences due to poor reads quality are denoted as X. Ensemble was used for the initial *NTNG* paralogs retrieval in marmoset (Ma) and mouse (Mo). Yellow (vertical strip) denotes the position of the SNP-related mutation allele and non-matched to human substitutions. (**E**) Evolution rates for the proteins encoded by the *NTNG7* (Netrin-G1) and *NTNG2* (Netrin-G2). (**F**) Amino acid changes for Netrin-Gs across hominins, primates and mice. For Netrin-G1, there are no common mutations among primates, hominins and modern human. Es contains 5 point mutations absent in other hominins; Chi contains 1 mutation outside the depicted area (A81S); all 4 mutations for marmoset are shown, and 10 more extra mutations for mouse are not shown. For Netrin-G2, all hominins except Ice (relative to hg 19) carry a T346A mutation (exon 5-located and known as rs4962173), also detectable in primates and mouse. Chi’s Netrin-G2 differs from all hominins by S371A mutation (exon 6 – nested) and present in marmoset as S371V. Non-matched to hg 1 9 amino acids are highlighted as pink. Refer to **SM** for the full alignments.

It can be concluded that both genes (as paralogs) contribute to the cognitive scoring produced upon the implemented IQ testing but in a cognitive domain-complementary manner. *NTNG1* contributes to the VC and PS while *NTNG2* for the WM and PO domain scores (Figure 1C).

The robust link observed between a single SNP and affected cognitive domain IQ score (Figure 1) can be explained by some global dramatic perturbations caused by the presence of a mutated allele and/or a functinal importance of its context-dependent positioning on the gene (Figure 2A and 3A). To determine a potential significance of the SNP alleles’ epistatic environment we compared the nucleotide (nu) sequence within the immediate vicinity of a SNP allele positioning (50 nu upstream and downstream) in mice, primates and modern human. We compared all 11 SNPs used for the IQ screening and plotted the identity percent as a function of distance from the mutated allele position (Figure 2B and 3B, see Supplementary Materials = SM). We found that the identity percent distribution over the analysed areas of ±50 nu is not uniform and displays a SNP allele position-centred dramatic evolutionary changes pointing towards a potential functional significance of the immediate vicinity of a SNP as short as ±10 nu and not further, referred from here and beyond as a Region Of Interest (ROI) for each specific SNP allele. We calculated the rates of evolutionary changes for each ROI as a percent identity change over the lapsed mln yrs of evolution (Figure 2C and 3C). Among the 6 *NTNG*1-located SNP ROIs three of them display accelerated rates of evolution from marmoset to chimpanzee (rs2218404, rs628117, rs96501) when compared to the mouse-marmoset rates, and rs2218404 (affecting VC in human subjects) additionally demonstrates an accelerated rate of evolution on the chimpanzee to human path (Figure 2C). As for the *NTNG2* located ROIs (Figure 3–B), rs1105684 is remarkably consistent at displaying high evolutionary rates around 0.8 and, together with rs2274855, both have identical rates at the mouse-marmoset and chimpanzee-human paths, but differ dramatically at the marmoset-chimpanzee point (0.8 vs 0, respectively). rs2274855 is the only *NTNG2*-nested SNP ROI that underwent an AE from chimpanzee to human. Next we compared the DNA sequences of all 11 ROIs across mice, primates and extinct hominins *NTNG* gene paralogs (see ST2 for the datasets sources used for the genes reconstruction). T-allele of rs2218404 is detectable in marmoset and its position in mouse corresponds to adenosine (Figure 2D). G-alelle (associated with a higher VC score comparing to the human T-allele carriers) is found in Mesolithic hunter Loschbour (8,000 BC) but not in another ancient hunter Motala12 and is also present in other two hominins belonging to the Neolithic period, Iceman and Eskimos (5,300 and 4,000 yrs BC, respectively). rs628117 is the only *NTNG1*-related mutation near vicinity of which (±50 nu) is located an intra-hominins (Es, Ice, Lo) mutation (Figure 2B, low left). The next, PS-affecting, ROI of rs96501 displays an intricate path of T-allele evolution (associated with a higher PS score) being anciently conserved from mice to primates but later substituted on the less efficient (in terms of the generated IQ scores) C allele in Neanderthals, later again replaced by the T allele in Mesolithic hunters and coming back during the Neolithic times (Figure 2D). The first *NTNG2* SNP rs1105684 is located at the beginning of the gene and affects WM in healthy human subjects (Figure 3A). The origin of the T-allele is evolutionary bound to marmoset since its position in mice is occupied by another pyrimidine base C (Figure 3D). Next on the gene are two SNP alleles for rs7851893 and rs3824574 which do not affect IQ and similarly to rs1105684 are surrounded by highly conserved ROIs not only in hominins and chimpansee (100% identity) but also in marmoset (except 1 mutation for rs7851893). rs2149171 ROI (affecting the WM and PO scores) similarly to rs3824574 is 100% conserved across the all species (including the mouse) except the allele itself. The attenuating IQ C-allele position is occupied in mice genome by T but present in Iceman and Eskimos genes. A distinct evolutionary path is taken by another cognitive endophenotype-associated and affecting WM and PO scores A-allele of rs2274855 and its ROI (Figure 3D). Its position in mice is likely to be occupied by the C pyrimidine base (the software places a blank instead of it) which is gradually substituted on purine G in chimpanzee and misplaced by the lower IQ score-associated A-allele in Mesolithic hominins. And 20 nu downstream of the centre of ROI is located a modern human-specific point mutation translated into the T346A protein substitution (as described below).

**Figure 3.**
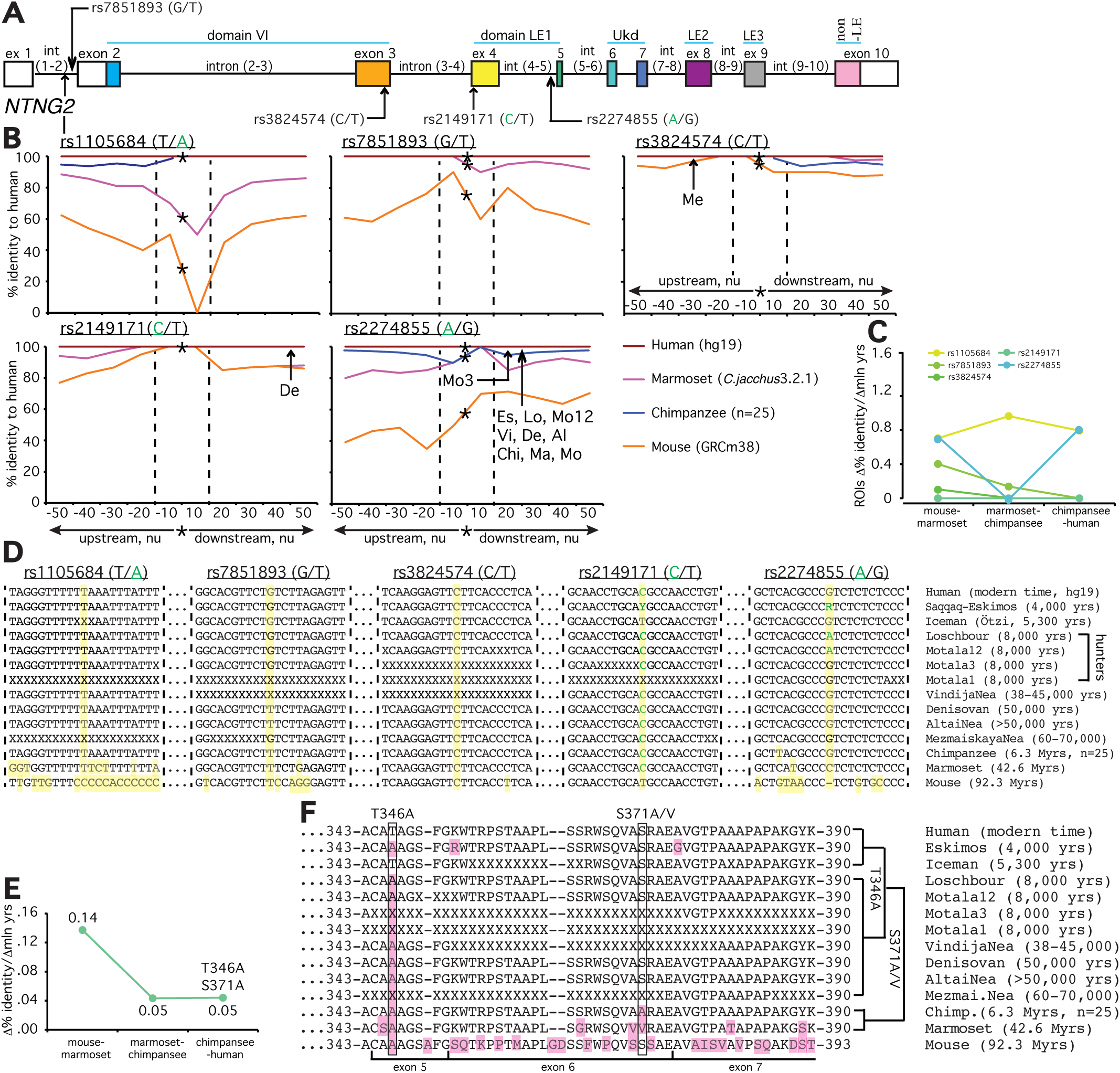
**Accelerated evolution (AE) and definition of a ROI of *NTNG* loci embedding associated with the cognitive endophenotype alleles. Hominins and primate-specific protein amino acid substitutions**. (**A**) *NTNG7* and *NTNG2* SNPs’ gene locations. Alleles associated with the lower IQ scores are shown either in red (*NTNG 7)* or green (*NTNG2).* (**B**) Calculated identity percent of primates and mice to human gene loci as a function of distance from the position of a mutated allele (denoted as a star). Comparison was done in a stepwise manner as +10 nu to the maximum of +50 nu using Stretcher (http://www.ebi.ac.uk/Tools/psa(**70**)). The areas were compared based on the positioning relative to the point mutation without any manual curation for “the best-fit” alignment, but as per the algorithm output only. Due to low level of identity for mouse and marmoset the initial search of the mutation allele positioning was done aligning against the human corresponding full-length intron, and then second time against the obtained 101 nu query (“-50 nu-SNP +50 nu”). Two dashed lines define an area of −10 nu to +10 nu from the mutation allele position. This area (21 nu in total) is defined as a ROI of the given mutation allele of a representing SNP. An arrow indicates a position of extinct hominins-specific mutations (see below). (**C**) Evolutionary rates for the ROIs calculated as a percent identity change relative to the hg 1 9 over the mln of years of evolution. The spectrum color reflects the mutations’ positioning order on a gene, as purple-yellow for *NTNG7,* and yellow-blue for *NTNG2.* (**D**) ROIs DNA sequences across hominins, primates and mice loci. The extinct and ancient hominin’s *NTNG* paralogs were reconstructed from the available datasets (see **SM**): Saqqaq-Eskimos (Es: **59**); Iceman (Ice: **58**); Hunters (Loschbour (Lo), Motala12 (Mo12), Motala3 (Mo3), Motala1: **57**); VindijaNea (Vi: **72**); Denisovan (De), AltaiNea (Al) and MezmaiskayaNea (Me): **73**; chimpanzee (Chi: reconstructed from (71), n=25 animals). Non-available sequences due to poor reads quality are denoted as X. Ensemble was used for the initial *NTNG* paralogs retrieval in marmoset (Ma) and mouse (Mo). Yellow (vertical strip) denotes the position of the SNP-related mutation allele and non-matched to human substitutions. (**E**) Evolution rates for the proteins encoded by the *NTNG7* (Netrin-G1) and *NTNG2* (Netrin-G2). (**F**) Amino acid changes for Netrin-Gs across hominins, primates and mice. For Netrin-G1, there are no common mutations among primates, hominins and modern human. Es contains 5 point mutations absent in other hominins; Chi contains 1 mutation outside the depicted area (A81S); all 4 mutations for marmoset are shown, and 10 more extra mutations for mouse are not shown. For Netrin-G2, all hominins except Ice (relative to hg 19) carry a T346A mutation (exon 5-located and known as rs4962173), also detectable in primates and mouse. Chi’s Netrin-G2 differs from all hominins by S371A mutation (exon 6 – nested) and present in marmoset as S371V. Non-matched to hg 1 9 amino acids are highlighted as pink. Refer to **SM** for the full alignments.

The distinct picture of evolutionary changes among the *NTNG1* and *NTNG2* nested SNPs has prompted us to compare evolutionary rates for the full-length proteins encoded by these gene paralogs, Netrin-G1 and Netrin-G2, respectively (2, 35). Netrin-G1 undergoes only few changes in its amino acid composition with the maximum calculated rate of evolution reaching 0.05 when mice and marmoset proteins are compared, 0.01 among the primates, and 0.03 between chimpanzee and human due to a single point mutation A81S (Figure 2E and SM: Netrin-G1), absent in other primates. Netrin-G2 evolves 2.8 times faster between mouse and marmoset than its paralog, Netrin-G1, and continues evolving with a steady rate of 0.05 from primates to human (Figure 3E and SM: Netrin-G2). We have also reconstructed both proteins from ancient (Neanderthals, Paleolithic time) and extinct hominins (Mesolithic and Neolithic times) and compared them with primates’ and mice’ Netrin-G orthologs (Figure 2F and 3F). Netrin-G1 is a highly conserved protein among the primates and hominins (Figure 3F). As for Netrin-G2, a mutation shared among the Neanderthals’ and Mesolithic genomes, primates and mice (T346A) is absent in the Neolithic Iceman and modern human (the signal for Motala3, Motala1 and MezmayskayaNea is not clear due to low sequence coverage). Primates (marmoset and chimpanzee) share another mutation (S371A/V) preserved in mouse and absent in hominins (further details can be found in the SM: Results).

## DISCUSSION

*NTNG* paralog SNPs and associated cognitive endophenotypes of human subjects. Shortcomings of cognitive and information processing are key features of SCZ diagnosis (36). They are frequently manifested as impairments in PO, WM, VC and PS (see (37) for references) and reported as attenuated scores upon IQ tests implementation. SCZ patients carrying a mutation allele for one of *NTNG* gene paralog SNPs form cognitive endophenotypes affecting the IQ scores (Figure 1C). The term “endophenotype” was originally coined by (38) and later advanced through the field of psychiatry by Gottesman and Shields (reviewed in (39)) as a biomarker associated with a specific phenotypic trait (40). The formed SCZ endophenotypic groups comprise from subjects with either affected VIQ (via attenuated VC by *NTNG1* rs2218404 or WM by *NTNG2* rs2149171 and rs2274855) or affected PIQ (via attenuated PO by *NTNG2* rs2149171 and rs2274855). In two extra cases PS is affected by rs96501 of *NTNG1* and WM by rs1105684 of *NTNG2* (Figure 1B-1 and B–2, respectively) but in healthy human subjects. Such intriguing non-overlapping effect on the IQ domains prompts us to conclude that *NTNG* paralogs complement each other function and represent an example of how a synapse-expressed genes affect the human cognitive abilities, perhaps through the precision of neuronal connectivity perturbations and concomitant miswirings. The observed phenomena of the affected WM is the most striking due to its multifaceted constructive nature (41) underlying many, if not all, cognitive tasks such as comprehension, reasoning and learning (42) and historically introduced by Baddeley as the reading span test (43). Lack of the localization effect of *NTNG2* SNP mutation alleles, all three are located in different parts of the gene (Figure 1A–2) but associated with identical endophenotype (Figure 1B–2), points to a uniform nature of the *NTNG2* function distribution over the entire gene. An obviously non-coding nature of all five IQ-affecting alleles (rs2149171 despite being exon 4-located encodes a silent F246F mutation) corroborates an idea that anthropoid trait-associated loci lie outside coding protein areas (44, 45). This hints towards a potential of these alleles to perturb genes regulatory functions, e.g. mRNA splicing, affecting downstream located pivotally functional *NTNG* elements such as Ukd-domain encoding exons 6 and 7 or a unique Netrin-Gs trait – GPI-link. Alternatively, or simultaneously, the *NTNG* SNP alleles may be embedded into an epistatic network of other genes influencing human cognitive traits (46). However since it is usual for a SNP effect to be estimated using an additive model (assuming either independent and cumulative single contribution) to the mean of a trait with the small effect size the power to detect the epistatic environment drastically declines. Contrary to the genetic associations with gene expression having large effect sizes (46), cognitive trait-associated effect sizes are reportedly small (47), e.g. the largest effect sizes of the variance of intelligence scores accounted for only 0.2% (48), 0.5% on GWA studies of 1,583 adolescence (49) or was predicted to be ~1% on 3,511 adults (50). Another GWAS of educational attainment (sharing a moderate correlate with intelligence), which included 126,559 individuals, reports on just 1% of the variance but only 0.02% in a replication sample (51). Our data support the preexisted conclusion that human cognitive traits modalities are not described by statistically large effect sizes.

Evolutionary elaborations of the embedding IQ-affecting mutations loci. Eleven previously published SCZ-associated SNPs were tested for their effect on IQ performance of human subjects and 5 of them were found to be associated with attenuated IQ cognitive endophenotypes (Figure 1B–1 and B–2). ROIs of 3 of them (rs2218404, rs1373336 and rs2274855) underwent an AE from chimpanzee to human when compared to New World monkeys to apes (marmoset-chimpanzee) path (Figure 2C and 3C). Two of them affect IQ in humans (rs2218404 – VC and rs2274855 – WM, Figure 1B–1 and B–2) while being located within the vicinity of exon 5 (2,275 nu downstream and 15 nu upstream, respectively) – a part of the lowest percent identity coding DNA among the *NTNG* gene paralogs (52).Presence of the evolutionary accelerated regions within the *NTNG* genes non-coding areas underscores them as contributors to the human-specific traits along with other genes (53). However, not only ROIs of the IQ-affecting alleles but the alleles themselves demonstrate several unique evolutionary features (see SM: Discussion). To understand evolutionary forces driving the emergence of cognitive endophenotype-associated alleles we have deduced a set of rules outlined as follows. 1. An alternative (mutated) allele evolutionary appearance coincides with the lack of any other mutations within ROI (a conserved island rule); 2. positioning of the future mutation often represents a turning point of dramatic changes of an allele ROI (e.g. as seen in marmoset: rs2218404 (50-90%), rs628117 (30-80%), rs96501 (100-40%)); 3. an AE of ROI often precedes the emergence of a mutation allele (e.g. rs2218404: chimpanzee to human (*k* = 1.59), appearance of “G” in Loschbour; rs628117: marmoset to chimpanzee (*k* = 1.10), appearance of “T” in AltaiNea; rs96501: marmoset to chimpanzee (*k* = 0.83), appearance of “C” in AltaiNea; rs2274855: chimpanzee to human (*k* = 0.79), appearance of “A” in Motala12; 4. low identity percent (equivalent to subsequent substantial evolutionary changes) among the evolutionary species within the allele surrounding proximity of as long as ±50 nu is not sufficient for the future mutated allele significance as a cognitive endophenotype determinant (as deduced by the IQ score) as seen for the rs1373336, rs1444042, rs4915045 and rs7851893 (none of them are IQ-affecting, though associated with SCZ, despite showing (very) low identity in mice). Rather some dramatic changes within the allele’s immediate proximity of ±10 nu (defined as a ROI) preceded or followed by more stringently conserved DNA are necessary (the conserved island perturbation rule). Currently we are unable to state that the IQ-associated alleles ROIs represent regulatory loci and an important source of evolutionary innovation (54) but they may be the smallest functional blocks of a strong positive selection that exerts its action upon, similarly to the 20–30 nu clusters of strongly conserved non-coding elements (CNEs),transcription factor binding sites (TFBS), RNA splicing and editing motifs (55).

Extinct hominins and IQ-associated mutation alleles. Availability of archaic genomes allows excavation for the advantageous alleles that modern humans acquired from archaic extinct hominins such as Neandertals and Denisovans who used to live 230,000-30,000 years ago (Middle/Upper Paleolithic, Old Stone Age) defined by distinct morphological features (56), and from modern extinct humans (hunters, farmers) from Mesolithic (Middle Stone Age, ~10,000 yrs BC, (57)) and Neolithic (New Stone Age, ~5,000 BC, (58, 59) periods. Though exhibiting several anatomical features, making archaic hominins different from the modern human, there are studies challenging the idea that reserve symbolism and abstract thinking was an exclusive prerogative of modern human (60, 61). The time Neanderthals used to live in is thought to be associated with the onset of cognitive fluidity involving the capacity to draw analogies (early paintings), to combine concepts (making tools) and to adapt ideas for new contexts (62). Wynn and Coolidge believe that evolution of WM was central to the evolution of human cognitive traits consisted from few genetic mutations that led to “enhanced WM” 200,000-40,000 BC (63). Our work partially supports this idea showing the perseverance of higher WM score-associated alleles across the hominins such as T of rs1105684 and G of rs2274855 (Figure 3D) but a Neolithic appearance of rs2149171 T in Iceman previously found only in mice genome supporting the conclusion made by Crabtree (64) that modern humans as species “are surprisingly intellectually fragile and perhaps reached a peak 2,000–6,000 years ago”. Mesolithic period has been always considered as a key gate for the evolution of human languages (65) with our data showing that rs2218404 G-allele associated with a higher VC score (Figure 1B–1) emerges for the first time in the Loschbour hunter *NTNG1* gene (Figure 3D). VC, as an example of abstract symbol usage, is associated with the global network efficiency (as a part of fronto-parietal network (19,66), and global communication and intellectual performance (67). From this point of view it is not surprising that appearance of the G allele in Loschbour and Iceman genomes coincides with the presence of PS enhancing rs96501 T allele (Figure 3D). A wealth of data has been collected characterising possible look and health status of archaic hominins and modern but extinct humans (for ref. see (68) and SM: Discussion). Based on our own data we may also speculate that the extinct hominins may have had a lower VC comparing to us, and consequently, Neanderthals were unlikely able for a semantic communication due to a global network inefficiency VC is associated with; they have had similar to us PS (if they had lived beyond the Mesolithic period), and were likely to have had identical to modern human WM, corroborating the advanced evolutionary nature of this important human cognitive domain of a limited capacity and associated with intelligence.

## CONCLUSION

Evolution of a novel function relies on enhanced genetic robustness via functional redundancy potentially provided by a gene duplication event. Further evolutionary outcome depends on the substrate availability (undergoing its own evolution) upon which the novel function(s) exerts its action. Nature does not create but tinkers to perfection provided to it material exploring available evolutionary tools. Half a billion years ago a gene duplication event had provided a plethora of such substrate thus converting the evolution itself into a “Creator” of new functions. Here we have described how a pair of twin genes got themselves involved into the human cognitive functioning believed to be emerged in a primordial state in primitive vertebrates prior to the first recorded gene duplication. Subsequent process of the function specialisation made *NTNG* paralogs to subfunctionalise into distinct cognitive domains in a complementary manner (52).

### MATERIALS AND METHODS

Ethics statement. This study was performed in accordance with the World Medical Association’s Declaration of Helsinki and approved by the Osaka University Research Ethics Committee. A written informed consent was obtained from all subjects after the procedures had been fully explained.

Subjects. The procedures were performed as per established protocols at Osaka University as described previously (69). The subjects consisted from 339 patients with SCZ and 716 healthy controls. The sex ratio did not differ significantly between the groups, but the mean age was significantly different. The subjects were all biologically unrelated Japanese and recruited from both outpatient and inpatient units at Osaka University Hospital and other psychiatric hospitals. Each patient with SCZ had been diagnosed by at least two trained psychiatrists based on unstructured clinical interviews, according to the criteria of the DSM-IV (36). In case if the diagnosis of the two trained psychiatrists was discordant, it was resolved through the further negotiations on both specialist opinions. In case of unresolved diagnostic disputes, the patient was omitted from the study. Psychiatrically healthy controls were recruited through local advertisements and were evaluated by means of unstructured interviews to exclude individuals with current or past contact with psychiatric services, those who experienced psychiatric medications, or who were not Japanese. Controls for family history of a CD, such as SCZ, BD, or major depressive disorder were not included. Ethnicity was determined by self-report and was not confirmed by genetic analyses. Additionally, subjects were excluded from this study if they had neurologic or medical conditions that could have potentially affect their central nervous system, such as atypical headaches, head trauma with loss of consciousness, chronic lung disease, kidney disease, chronic hepatic disease, thyroid disease, active cancer, cerebrovascular disease, epilepsy, seizures, substance abuse related disorders, or mental retardation.

SNPs selection, genotyping, and genomic sequencing. This study was designed to examine the association of SCZ patients cognitive performance (through WAIS-III implementation (17)) with *NTNG* genes. Venous blood was collected from the subjects. Genomic DNA was extracted from the whole blood using standard procedures. The SNPs (26–28, 30) were genotyped using the TaqMan allelic discrimination assay (Applied Biosystems, Foster City, CA). No deviations from the Hardy-Weinberg equilibrium in the examined SNPs were detected (*p* > 0.05).

Statistical analysis. The effects of the diagnosis, genotype and their interaction on cognitive performances in the WAIS were analyzed by two-way analyses of covariance (ANCOVA). Diagnosis and genotype statuses were included in the model as independent variables (ST1). FIQ and each WAIS subscale score (VIQ, PIQ, VC, PO, WM, PS, Vc, SiM, IF, CH, AM, DS, LNS, PC, BD, and MR) were included as dependent variables. Sex, age and years of education were treated as covariates, as they were possible confounding factors. All *p* values are two tailed, and statistical significance was defined as *p < 0.05 and **p < 0.01.

Identity percent calculations and the definition of ROIs. The complete procedure is described in the Figures 2–3 legend. Stretcher (70) was used for the alignments (the default values were: gap penalty – 16 (DNA) and 12 (protein), and the extend penalty – 4 (DNA) and 2 (protein)), for the percent identity calculations and evolutionary rates. A ROI’s selection was initially defined as a minimal area surrounding a SNP mutation allele incorporating the outmost evolutionary dramatic changes, turns out to be ±10 nu.

Mice, primates and hominins *NTNG* paralogs DNA and encoded protein sequences reconstruction. Genomes for mouse (GRC38.p3) and marmoset (C_jacchus3.2.1) were from Ensemble. Since chimpanzee’s genome is based only on a single individual (CHIMP2.1.4, Clint) and contains several questionable information we have reconstructed a consensus genome sequence for both *NTNG* genes based on 25 primate sequences of *Pan troglodytes* (71). All datasets used for the *NTNG* paralogs DNA and encoded by them proteins reconstruction are listed in ST2. For details refer to SM.

## SUPPLEMENTARY MATERIALS (SM)

Contain additional Results and Discussion, Supplementary Methods (ancient and primate genomes reconstructions), and Supplementary Tables (ST1 and ST2) as a single compiled pdf file. Also included are human Netrin-G1 and Netrin-G2 alignments, as well as 101 nu alignments for all 11 ROIs across the all analysed species.

## ACKNOWLEDGEMENTS

Authors would like to acknowledge the financial support provided by Funding Program for World-Leading Innovative R&D on Science and Technology (FIRST Programme) and KAKENHI 15H04290 by the Japan Society for the Promotion of Science (JSPS).

